# Binding of cardiolipin to the KcsA channel at the membrane outer leaflet allosterically opens the inner gate

**DOI:** 10.1101/2022.02.08.479071

**Authors:** Masataka Inada, Masayuki Iwamoto, Norio Yoshida, Shigetoshi Oiki, Nobuaki Matsumori

## Abstract

Membrane proteins embedded in the membrane undergo changes in their actions under the influence of membrane lipids. Here, we present a novel type of lipid action on the potassium channel KcsA from *Streptomyces lividans*. Cardiolipin is present in various cellular membranes, including the host membrane of KcsA. Although the M0 domain, a nontransmembrane helix, is known to sense anionic lipids in the inner leaflet, we found that divalent anionic cardiolipin in the outer leaflet of the membrane interacts with positively charged residues, Arg64 and Arg89, on the extracellular side of the transmembrane domain. This binding propagates its action across the membrane toward the intracellular region of KcsA, thus, opening the inner gate. Such a long-range allosteric effect has not been found for channel–lipid interactions.

## Introduction

The structural and functional integrity of membrane proteins (MPs) is secured by their membrane environment (1-14). The effects of membranes on MP structure and function can be explained by 1) direct interactions between membrane lipids and MPs (2-10) and 2) membrane physicochemical properties, such as thickness, tension, and lateral pressure (11-16). In particular, recent crystallographic studies of MPs have revealed the presence of specifically binding lipid molecules (17-19), suggesting the frequent occurrence of direct MP–lipid interactions.

KcsA, from *Streptomyces lividans*, is a homotetrameric potassium channel with high structural similarity to eukaryotic congeners (20-22), and its high-resolution structure co-crystallized with lipid molecules has been resolved (22). KcsA is activated by intracellular acidic pH (23-26) (Fig. 1), but to be fully open, a specific lipid environment is a prerequisite: negatively charged phospholipids in the inner leaflet bind to the N-terminal amphipathic M0 helix, lying at the membrane interface when the channel is in the acid-activated state, stabilizing the open conformation (27, 28). This lipid-sensing M0-helix is, however, not a transmembrane helix, and most MPs do not have such sensors to explore the membrane environment. Generally, membrane lipids interact with transmembrane domains (TMDs) directly at their interface to the membrane phase (17-19); hence, it is logical to assume that the binding of lipids to the TMD of KcsA is also involved in the regulation of channel activity. Although various types of KcsA–lipid interactions have been found (28-31), systematic studies on the topic have not been performed, mainly because of the lack of a practical experimental methodology.

**Fig. 1.**
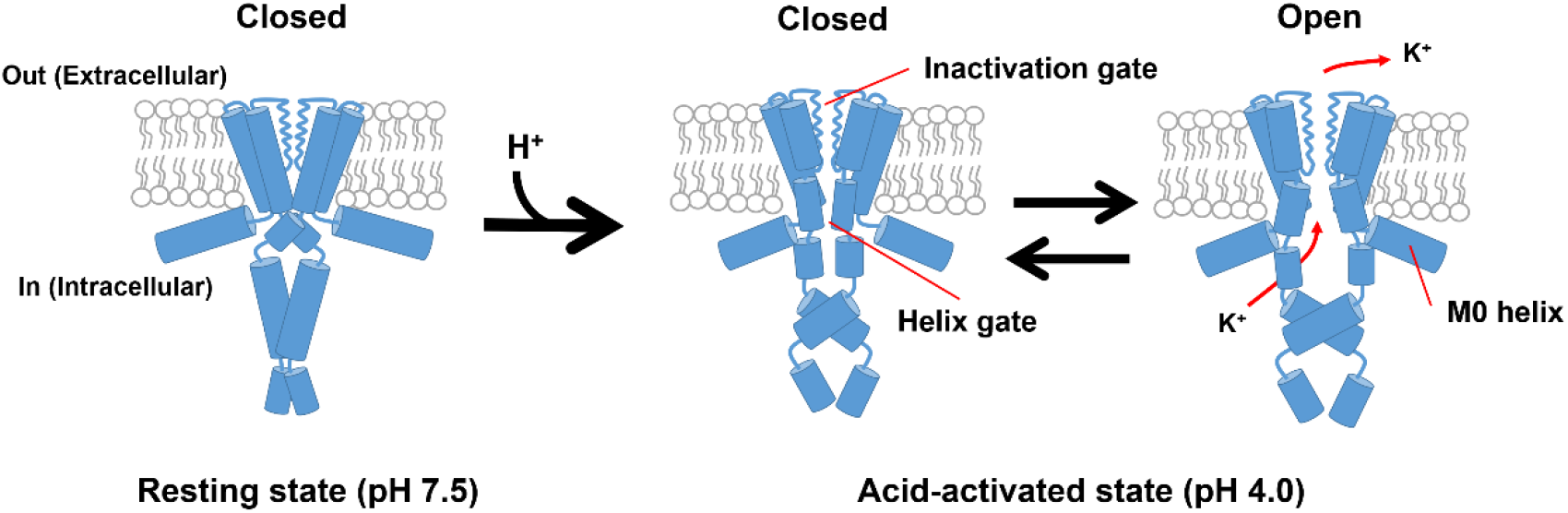
Gating states of the KcsA channel. The gate does not open at neutral pH. At acidic pH on the intracellular side, the channel becomes acid-activated; the pH sensor in the cytoplasmic domain changes conformation, and the inner gate undergoes open-close conformational changes via a global twisting motion.

We recently developed a surface plasmon resonance (SPR)-based method for the quantitative analysis of the interactions between MPs and lipids (Fig. 2A) (32), which was named SAMPLIA (Self-assembled monolayer-Assisted MP-Lipid Interaction Analysis). In this method, the sensor chip surface was modified with a self-assembled monolayer (SAM), which allows for a larger immobilization of MPs and provides a partial membranous environment to the immobilized MPs. The affinity of various types of lipids to MPs is sensitively and quantitatively evaluated by the addition of a solubilized lipid solution to the immobilized MPs. In this study, the SAMPLIA method was applied to quantitatively analyze the interactions between KcsA and anionic lipids. Although phosphatidylglycerol (PG) has been the most frequently used to test the effect of lipids on the KcsA channel (28-31), PG is not present in native *Streptomyces* membranes (33-35). Alternatively, phosphatidic acid (PA) and cardiolipin (CL) are major anionic lipid constituents of the *Streptomyces* membrane (33), and these lipids, in addition to PG as a positive control, were used in the present analysis. To gain insight into the binding site of these lipids, wild-type KcsA (wt-KcsA) and a deletion mutant lacking the anionic lipid-sensor domain, M0, as well as a non-inactivating mutant and mutants at Arg residues in the outer leaflet (extracellular) side were used for the interaction analysis. Single-channel recordings using the contact bubble bilayer (CBB) method were carried out to examine the functional contribution of these lipids to the gating of the KcsA channel. The combined approach with the SAMPLIA method and the CBB method revealed unprecedented actions of CL on the KcsA channel.

**Fig. 2.**
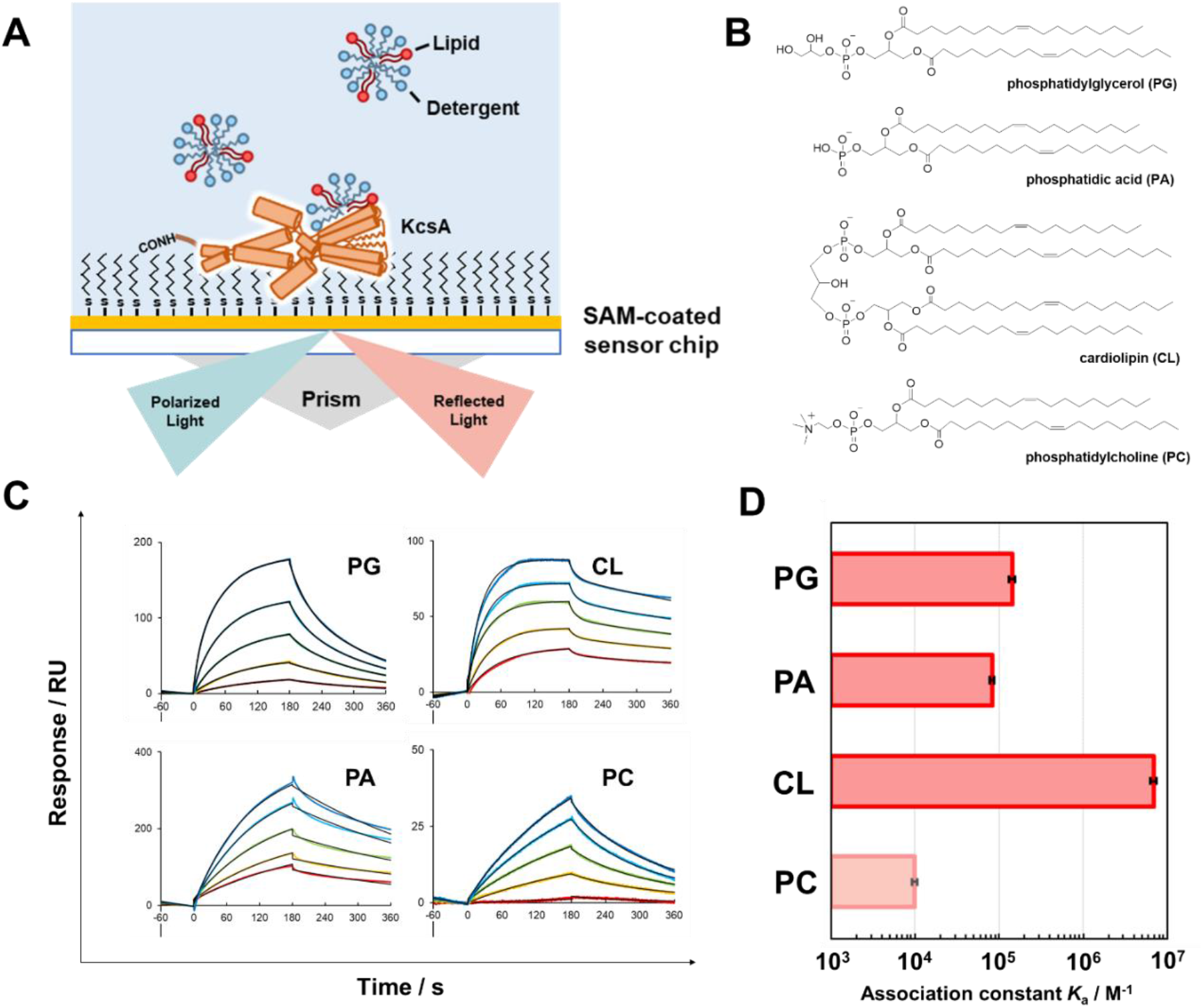
Analysis of the interaction of phosphatidylglycerol (PG), phosphatidic acid (PA), cardiolipin (CL), and phosphatidylcholine (PC) with wt-KcsA. **(A)** Schematic diagram of SAMPLIA method for KcsA–lipid interaction. KcsA molecules are covalently immobilized onto the sensor chip coated with SAM composed of mercaptocarboxylic acid. Solubilized lipids are added to detect the interaction with the MP (32). **(B)** Chemical structures of PG, PA, CL, and PC. **(C)** SPR sensorgrams showing the interactions of PG, PC, PA and CL with wt-KcsA immobilized on the C_6_-SAM modified sensor chip. 10 (bottom), 20, 30, 40, and 50 (top) μM of the lipids dissolved in acidic buffer [10 mM succinic acid, pH 4.0, 200 mM KCl, 3 mM EDTA, 0.05 %(v/v) Tween 20] were injected at a flow rate of 20 μL min^−1^. Black lines indicate experimentally obtained sensorgrams, while red lines indicate theoretical curves. **(D)** The affinity of PG, PA, CL, and PC toward wt-KcsA in acidic buffer (10 mM succinic acid [pH 4.0], 200 mM KCl, 3 mM EDTA), calculated from the sensorgrams by fitting to a 1:2 heterogeneous ligand binding model, as listed in Table S2, and higher association constants were adopted (see text).

## Results and Discussion

### CL Interacts with KcsA more strongly than PG and PA

KcsA was immobilized on the SAM-modified sensor chips at ∼3000 RU, which enabled us to clearly observe the interactions between MPs and lipid molecules (Fig. 2A). According to the crystal structure of KcsA (20, 22), the length of the SAM (ca. 1 nm) was not thick enough to completely cover and accommodate the TMDs of KcsA, and thus, a considerable part of the transmembrane helices should be outside of the SAM surface. This allows for unrestricted contact between the immobilized KcsA and the extraneously added lipids. We further evaluated the particle size of lipid-detergent mixtures and confirmed that the lipids were solubilized in the SPR running buffer containing the detergent Tween 20 (Table S1).

Prior to analyzing the interaction between wt-KcsA and lipids, we examined the interaction between immobilized wt-KcsA and tetrabutylammonium (TBA), a specific blocker of KcsA (Fig. S1A) (36). KcsA co-crystallized with TBA revealed a binding configuration of TBA with its four butyl chains extending to the subunits (37). The high-affinity binding of TBA to KcsA deduced from the present SPR study (Fig. S1B, C) strongly suggests that KcsA retains its tetrameric form on the chip (Fig. 2A) (36). Next, we evaluated the interaction between PG and wt-KcsA to confirm the validity of the SAMPLIA method (28-31). Since KcsA adopts an acid-activated state at acidic pH and a resting state at neutral pH (Fig. 1), both acidic (pH 4.0) and neutral (pH 7.5) buffers were used for the analysis. The observed sensorgrams were fitted by a 1:2 heterogeneous ligand binding model that assumes that the lipid binds to KcsA at two different binding cites (Table S2). Since the smaller association constants (*K*_A_) are considered to be due to non-specific or less relevant binding, because their maximum binding values (*R*_max_) are much smaller than those of the larger *K*_A_s (Table S2), indicating their little contribution to the binding. Therefore, we focused on the larger *K*_A_ values in the following experiments. The results revealed that wt-KcsA binds PG more strongly at pH 4.0, than at pH 7.5 (Fig. 2C, Fig. S2, Table S2), and the binding of PG to wt-KcsA was much stronger than that of phosphatidylcholine (PC) to wt-KcsA (Fig. 2C, D, Table S2). This result is consistent with those of previous studies that the KcsA channel opens in the presence of PG at acidic pH (28,30,31), thus, verifying that the SAMPLIA method can be used to properly evaluate the KcsA–lipid interactions.

We subsequently evaluated the interaction of wt-KcsA with PA and CL, both of which are major components of *Streptomyces* membranes. These lipids showed less significant interaction with resting state KcsA at pH 7.5 (Table S2); in particular, CL did not show any observable binding to KcsA at pH 7.5 (Fig. S2, Table S2). In contrast, at acidic pH, where KcsA adopts an acid-activated state, the interaction of CL with KcsA was stronger than those of PA and PG (Fig. 2, Table S2), suggesting that CL has the highest affinity for acid-activated KcsA.

### CL binds to KcsA at multiple sites including the M0 helix

Previous studies have shown that the M0 helix enables the sensing of anionic phospholipids in the membrane and stabilize the open conformation of KcsA (27, 28). To understand the contribution of the M0 helix to the high-affinity binding of CL, an M0 helix-deleted mutant of KcsA, ΔM0-KcsA (Fig. 3A), was used. The ΔM0-KcsA hardly influenced the affinity for CL, whereas it exhibited a significantly weakened interaction with PA and PG (Fig. 3B, C). These results indicate that CL has an additional binding site other than the M0 helix, while the M0 helix is the major binding site for PA and PG. The involvement of the M0 helix for CL binding was evaluated using an M0 helix-mimicking peptide, and CL bounds to the M0 helix (Fig. S3). These results indicate that CL interacts with acid-activated KcsA at multiple sites, including the M0 helix, while PA and PG bind exclusively to the M0 helix.

**Fig. 3.**
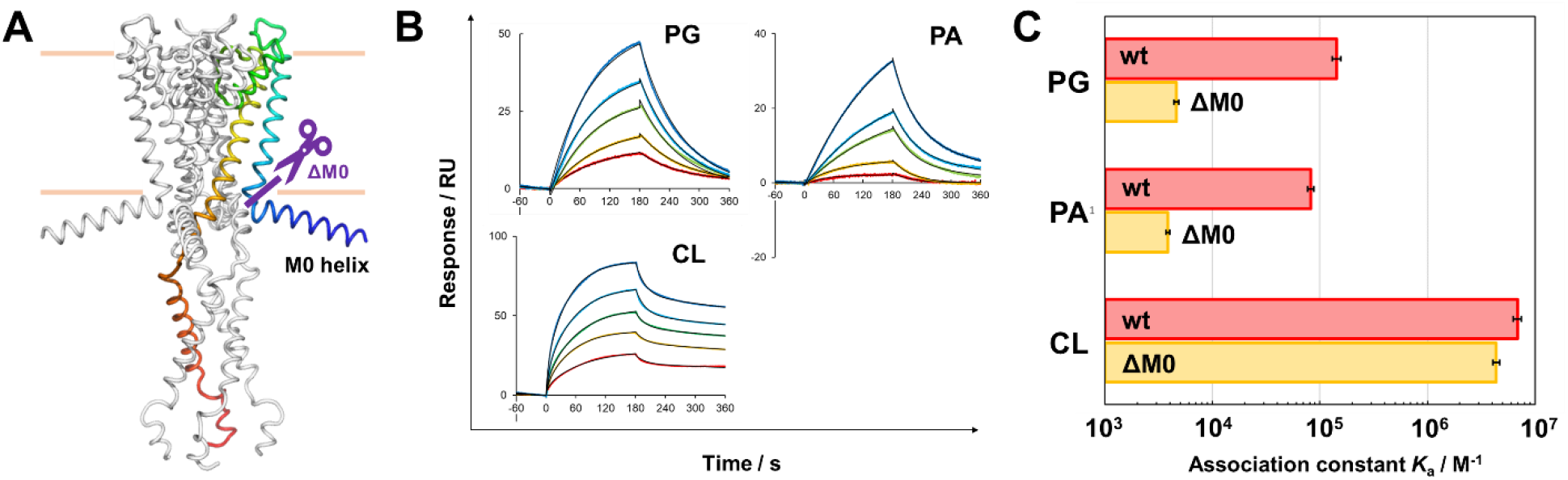
Effect of truncation of the M0 helix on the affinity of lipids toward KcsA. **(A)** The location of deletion on KcsA. ΔM0-KcsA is the mutant in which the N-terminal M0-helix domain was truncated (purple line). **(B)** SPR sensorgrams showing the interactions of PG, PA and CL with ΔM0-KcsA immobilized on the C6-SAM modified sensor chip. 10 (bottom), 20, 30, 40, and 50 (top) μM of the lipids dissolved in acidic buffer [10 mM succinic acid, pH 4.0, 200 mM KCl, 3 mM EDTA, 0.05 %(v/v) Tween 20] were injected at a flow rate of 20 μL min^−1^. Black lines indicate experimentally obtained sensorgrams, while red lines indicate theoretical curves. **(C)** The affinity of PG, PA, and CL toward wt-KcsA and ΔM0-KcsA in acidic buffer (pH 4.0). Each affinity was calculated from the sensorgram by fitting to a 1:2 heterogeneous ligand binding model, as listed in Table S3.

### Distinct mechanism of CL on the KcsA channel activity

To gain further insight into the distinct mode of CL action, we performed electrophysiological measurements using the contact bubble bilayer (CBB) method (Fig. 4A, B) (38), and assessed the effect of lipids on KcsA activity. The single-channel current of the acid-activated KcsA (Fig. 1) was recorded (Fig. 4B) (38-40). First, as a control experiment, the lipid bilayer was formed using pure 1-palmitoyl-2-oleoyl-sn-glycero-3-phosphocholine (POPC), and the open probability (*P*_open_) was approximately 1% (Fig. 4C, D). In the presence of anionic lipids (25 wt%), the channel exhibited substantially high *P*_open_ values, i.e., 17% for the PG-containing membrane (*In/Out* in Fig. 4C, D). These results are comparable to those of a previous report, in which the bilayer contained 100% anionic lipids (28). In the case of CL (*In/Out* in Fig. 4C, D), the *P*_open_ value (30%) was significantly higher than those of other anionic lipids (p < 0.05), which is relevant to the higher CL binding affinity revealed by the SPR experiments (Fig. 2).

**Fig. 4.**
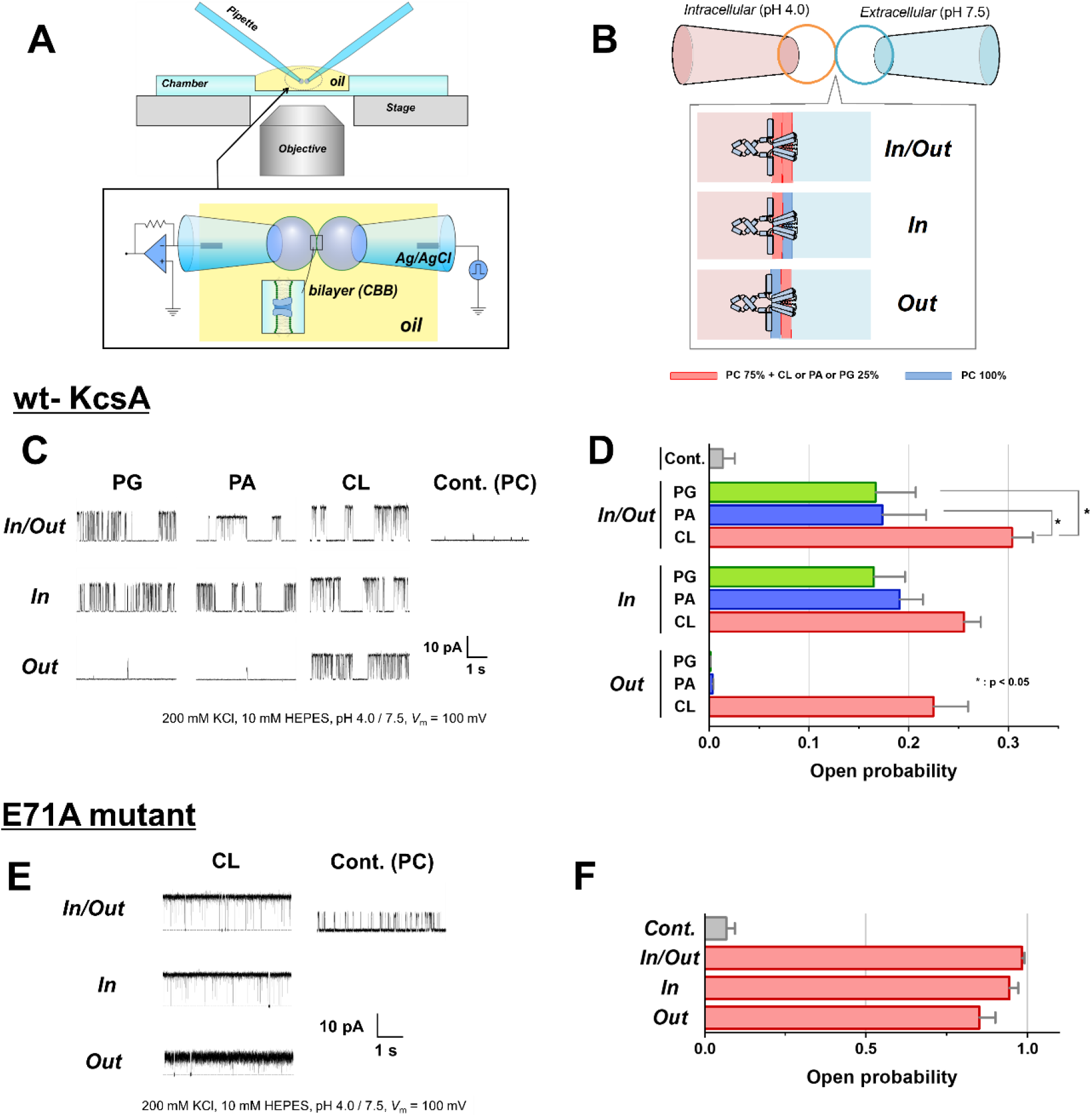
Single-channel recordings using the contact bubble bilayer (CBB) method. **(A)** The CBB method. Water bubbles are blown from two glass pipettes (left bubble: pH 4.0 for intracellular side; right bubble: PH 7.5 for extracellular side) into the oil (hexadecane). Two bubbles with their surface lined by a lipid monolayer contact for the bilayer formation. **(B)** Symmetric (In/Out) or asymmetric (In, Out) membrane and the orientation of KcsA in the CBB. The KcsA channel is activated by intracellular acidic pH. Therefore, by making the pH conditions asymmetric (pH4/pH7.5), the orientation of the “active” channel in the membrane can be controlled so that only channel molecule with its cytoplasmic domain facing acidic bubble contributes to the single-channel current. In the CBB experiment, the membrane leaflet facing the acidic bubble (left) is defined as the inner leaflet, and the other (right) is the outer leaflet. **(C)** Single-channel current traces of the wt-KcsA channel in symmetric and asymmetric membranes at 100 mV. Single-channel conductance differs substantially in different lipid membranes, owing to the surface potential of negatively charged lipids (28). **(D)** Open probability (*P*_open_) of the symmetric or asymmetric membrane with different compositions for wt-KcsA. Error bars represent SEM (n = 3-14). *P*_open_ values are listed in Table S5. **(E)** Single-channel current traces of the E71A mutant in symmetric and asymmetric membranes at 100 mV. **(F)** *P*_open_ of the E71A mutant in symmetric or asymmetric CL-containing membrane. Error bars represent SEM (n = 4-8). *P*_open_ values are listed in Table S6.

To explore the binding sites for these anionic lipids, single-channel recordings were performed in asymmetric membranes, in which one leaflet contained pure POPC and another leaflet contained 25% anionic lipids in POPC (*In* or *Out* in Fig. 4B). When the inner leaflet contained anionic lipids (*In* in Fig. 4C), the channel activity was similar to that in the symmetric membranes (*In/Out* in Fig. 4C), that is, the *P*_open_ values for the PG-, PA-, and CL-containing membranes (*In* in Fig. 4D) were indistinguishable from those in the symmetric membranes (*In/Out* in Fig. 4D). Thus, we confirmed that the KcsA channel was opened by anionic lipids in the inner leaflet (28, 41), which is consistent with the tight binding of the lipids to the M0 helix, as demonstrated by the SPR data using both M0-truncated KcsA (Fig. 3) and the M0 helix peptide (Fig. S3C).

Strikingly, we discovered that CL efficiently opened the KcsA channel from the outer leaflet (extracellular side of the TMD) (*Out* in Fig. 4C). The single-channel activity upon the presence of outer leaflet CL was indistinguishable from that when CL was contained in both sides or in the inner leaflet; *P*_open_ was more than 20% (Fig. 4D and Table S5). This is consistent with the SPR data showing that CL binds tightly to the M0-deleted mutant (Fig. 3). Accordingly, we demonstrate for the first time that CL binds to the outer (extracellular) half of the TMD and activates the KcsA channel.

To explore the mechanism underlying the action of CL in the outer leaflet, the E71A mutant was used. Previous reports using the non-inactivating mutant of E71A, in which the inactivation gate remains open, demonstrated that the M0-helix mediated action of anionic lipids opens the activation gate, or the inner gate (28, 42). The single-channel currents of the E71A mutant in the presence of CL in either or both leaflets are shown in Fig. 4E. The *P*_open_ was nearly 100% in all cases (Fig. 4F, Table S6), which unequivocally indicated that the outer leaflet CL opened the inner gate. The potential effects of the outer CL on the inactivation gate can be referred to by inspecting the wt-KcsA channel in Fig. 4D. In the presence of CL on either side or both sides, the *P*_open_ values were statistically indistinguishable, suggesting that the effect of CL on the inactivation gate was negligible. Taken together, our electrophysiological experiments revealed that CL existing in the outer leaflet membrane selectively opened the inner gate located in the intracellular half of the TMD. Therefore, CL in the outer leaflet functions as an allosteric modulator (10, 43).

### CL binds to Arg residues in the extracellular surface of KcsA

The SPR and CBB experiments demonstrated that CL binds not only to the M0 helix but also to the outer leaflet side of the protein and opens the inner gate. We then examined the CL binding site on the outer (extracellular) side of KcsA. As CL is a divalent anion with two phosphate groups near the membrane interface (Fig. 2B), we assumed that two positively charged residues (Arg64 and Arg89), both of which are located at the outer membrane interface of the TMD and face inter-subunit space (17, 44) (Fig. 5A), serve as counterparts for CL binding. To prove this, we examined the SAMPLIA and CBB analyses of R52Q, R64Q, and R89Q mutants. Mutations at Arg64 and/or Arg89 significantly reduced the affinity for CL (Fig. 5B, C); concurrently, the mutants abolished the gate opening induced by the outer leaflet CL (Fig. 5D, *Out*). In contrast, the mutation at Arg52, which is also located at the outer membrane interface of the TMD (Fig. 5A), had no effect on either the binding affinity to CL (Fig. 5C) or CL-induced gate opening (Fig. 5D, E). These findings verify that CL selectively interacts with Arg64 and Arg89 and opens the inner gate located in the intracellular half of the TMD via a long-range allosteric coupling.

**Fig. 5.**
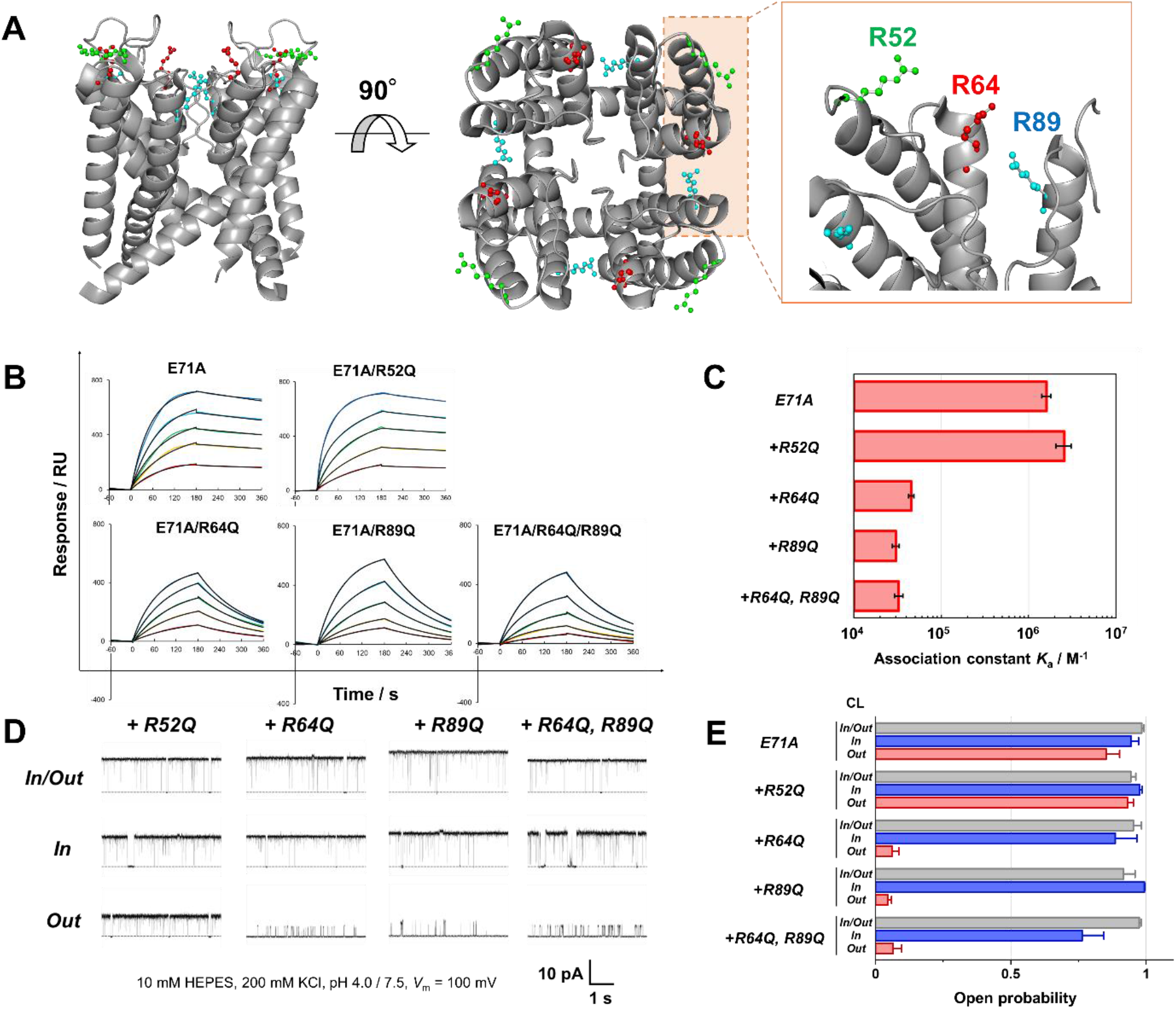
Effect of mutations of KcsA at Arg52, Arg64 and Arg89. (A) The positions of Arg52 (green), Arg64 (red) and Arg89 (blue) in KcsA (Protein Data Bank ID 6by3). All of the three residues are located at the outer membrane interface (extracellular side) of the TMD, while Arg64 and Arg89 also face the inter-subunit space. (B) SPR sensorgrams showing the interaction of CL with KcsA mutants, E71A, E71A/R52Q, E71A/R64Q, E71A/R89Q, and E71A/R64Q/R89Q, immobilized on the C6-SAM modified sensor chip. 10 (bottom), 20, 30, 40, and 50 (top) μM CL, solubilized in acidic buffer [10 mM succinic acid, pH 4.0, 200 mM KCl, 3 mM EDTA, 0.05 %(v/v) Tween 20], were injected at a flow rate of 20 μL min^−1^. Black lines indicate theoretical curves. (C) The affinity of CL to the KcsA mutants evaluated by the SAMPLIA method. Each affinity was calculated from the sensorgram in acidic buffer (pH 4.0) by fitting to a 1:2 heterogeneous ligand binding model, as listed in Table S7. (D) Single-channel current traces of the E71A/R52Q, E71A/R64Q, E71A/R89Q, E71A/R64Q/R89Q KcsA channel in symmetric and asymmetric membranes at 100 mV. The traces of E71A are shown in Fig. 4E. (E) *P*_open_ of the KcsA mutants in symmetric or asymmetric CL-containing membrane determined by the CBB method. Error bars represent SEM (n = 3-19). *P*_open_ values are listed in Table S8.

To gain further insight into the mode of interaction between CL and KcsA, we performed molecular dynamics (MD) simulations for KcsA embedded in POPC bilayers with various anionic lipids. The initial structure of KcsA was obtained from Protein Data Bank 6by3 (45), and the binding of PA, PG, and CL to the outer (extracellular) half of the respective KcsA channel was simulated (Fig. 6, Fig. S4). The radial distribution functions between the side chain N**H** atoms of Arg64 or Arg89 and the O atoms of anionic lipids showed that the position of the 1st peak was comparable for all lipids, but CL showed a slightly higher value (Fig. 6B). More importantly, CL tends to simultaneously form hydrogen bonds at both Arg64 and Arg89 (Fig. 6C). In fact, the time evolution of the number of hydrogen bonds shows that CL forms multiple hydrogen bonds more frequently than PA and PG (Fig. 6D, Fig. S4). The formation of such bridging structures may contribute to the higher affinity of CL for the outer half of KcsA as compared to the other anionic lipids.

**Fig. 6.**
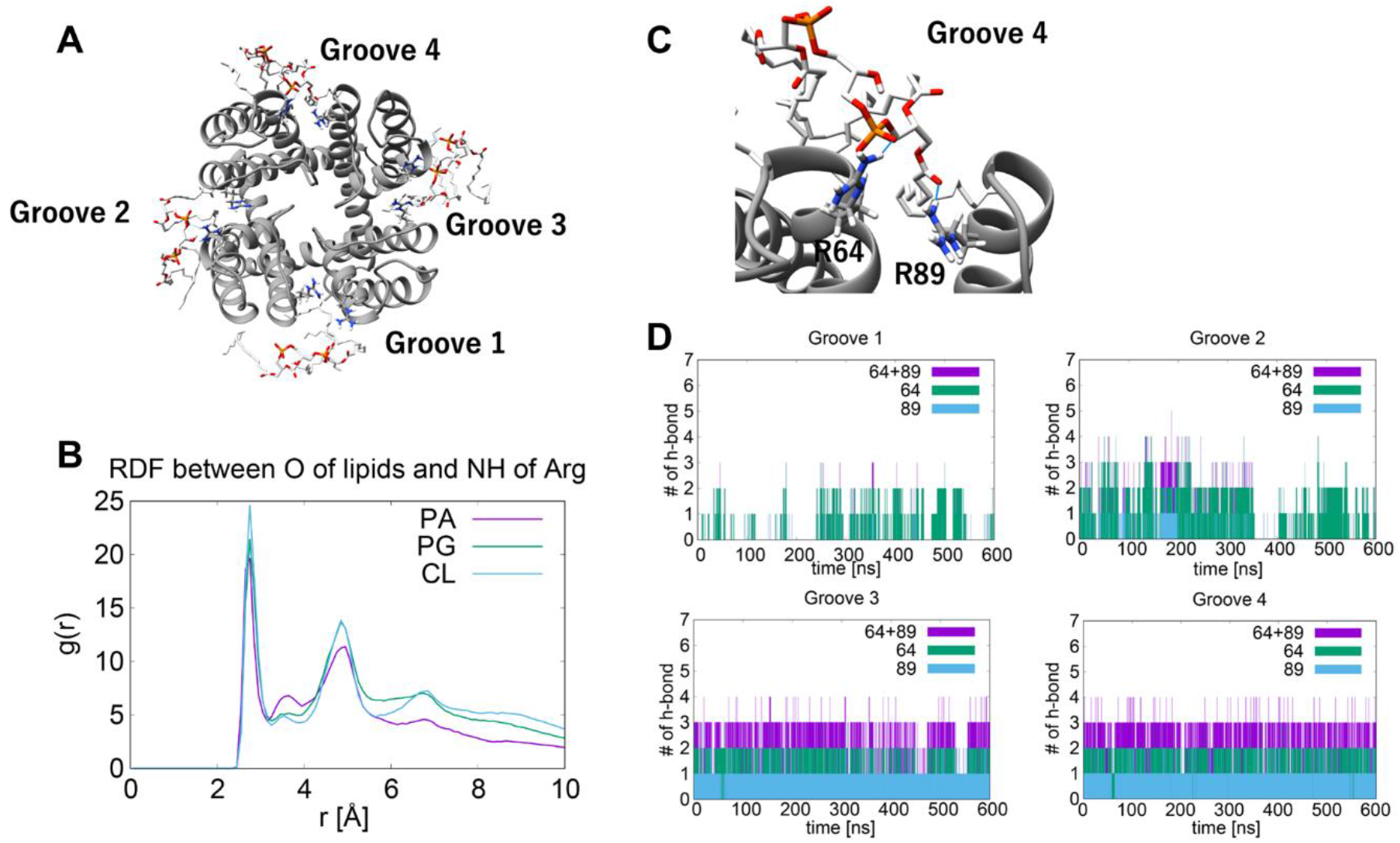
MD simulations for the binding of PA, PG and CL to KcsA. **(A)** Final snapshot of KcsA with the 1st neighbor CL to Arg64 and Arg89. **(B)** The radial distribution functions between side chain N**H** atoms of Arg64 or Arg89 and O atoms of anionic lipids. **(C)** Close-up view of the groove 4. Blue lines indicate the possible hydrogen-bond between CL and Arg64 or Arg89. **(D)** The number of hydrogen bond between CL and Arg64 or Arg89 at each groove. Purple traces indicate hydrogen bonds of CL with both Arg64 and Arg89. CL can simultaneously form hydrogen bonds with Arg64 and Arg89 to bridge them (see also Fig. S4).

Generally, potassium channels undergo conformational changes in the inner gate by undergoing a global twisting motion at acidic pH (Fig. 1) (23-26). Although the crystal structure of the closed and open conformations indicates that the extracellular half of the TMD is nearly unaltered in its conformation, the vigorous conformational change of the inner gate upon the channel opening (46) should induce a collective motion of the outer TMD (25). CL is likely to bind to Arg64 and Arg89 at the outer membrane interface of the TMD more readily in the open conformation and shift the equilibrium to the open state (Fig. 7), thus exerting its allostery. Such conformation-dependent binding of CL is partly supported by our finding that CL bound highly selectively to acid-activated KcsA (pH 4.0) compared to resting-state KcsA (pH 7.5), while the selectivity of PA and PG toward the acid-activated state was not so high (Fig. S2, Table S2).

**Fig. 7.**
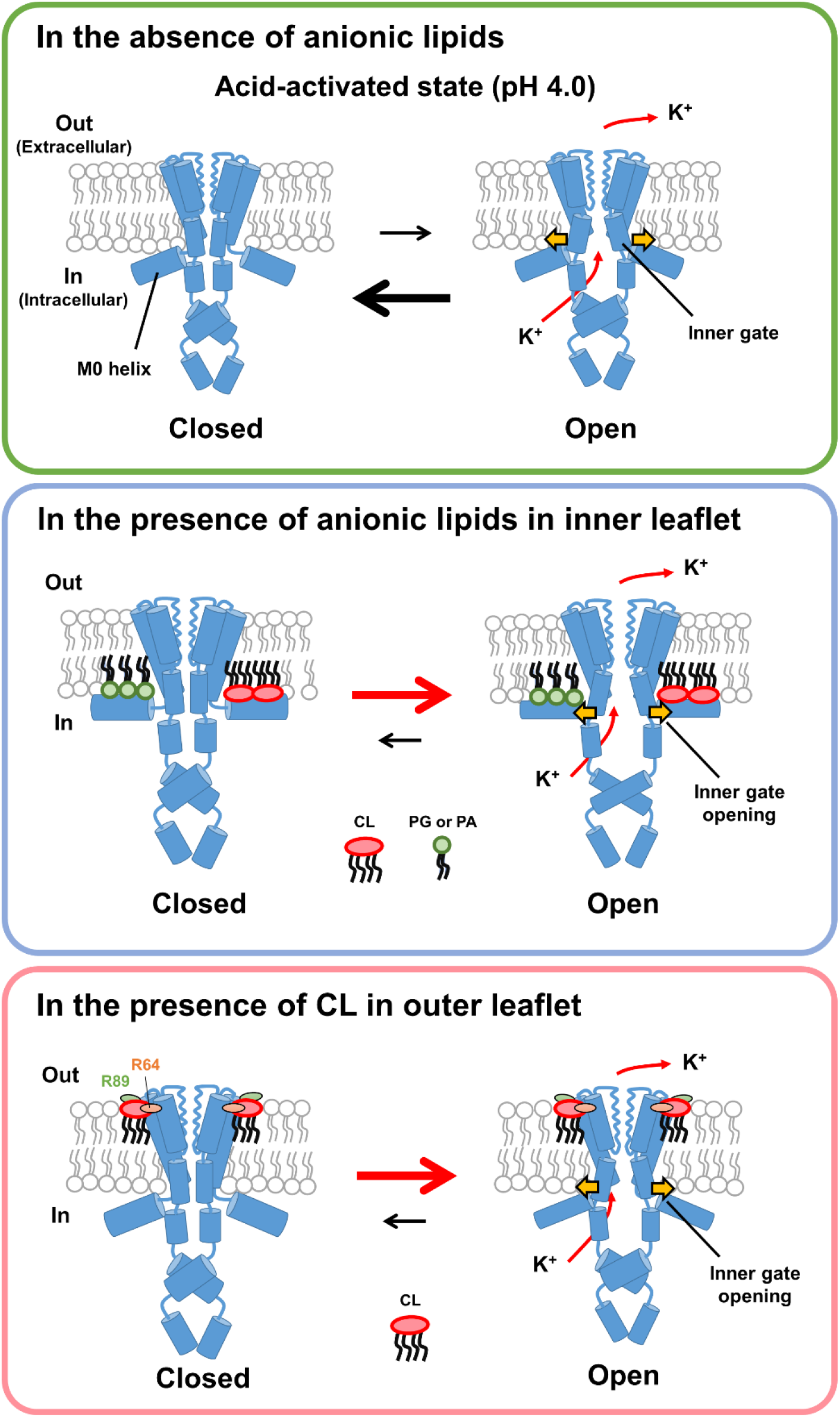
Model of the specific binding of anionic lipids to the KcsA channel and its effect on the inner gate via the two pathways. Upper panel: In the absence of anionic lipids, the equilibrium is inclined toward the closed state even in the acid-activated state. Middle panel: The M0 helix-mediates opening of the inner gate. All the anionic lipids examined (PA, CL, and PG) bind to the M0 helix tightly for the acid-activated channel (41). Bottom panel: Binding of CL to Arg64 and Arg89 opens the inner gate via an allosteric modulatory effect.

The KcsA channel has been used as a prototypical channel due to the extensive structural and functional information available, while the physiologically relevant function of the KcsA channel in *Streptomyces* native membranes has not been addressed. Unlike *E. coli* used for the expression of KcsA, *Streptomyces* does not possess PG in its membrane, and CL is a major component (33). Tight binding of CL to KcsA and the resulting channel activation provide insights into the physiological function of the KcsA channel *in situ* in the life cycle of *Streptomyces*.

## Conclusion

KcsA is a homotetrameric potassium channel containing the inner gate which opens under intracellular acidic pH (23-26). The conformational transition of the inner gate is shifted to an open conformation in the presence of anionic lipids in the membrane (Fig. 7) (41). The current study revealed that CL in the inner leaflet activates the KcsA channel by binding to the M0 peptide, as in the case of other anionic lipids (Fig. S3C, Table S4). Moreover, the high-affinity binding of CL to the M0-deleted mutant indicated the presence of an additional binding site for CL (Fig. 3). Active gating in the presence of CL in the outer leaflet (*Out* in Fig. 4C, D) indicates that CL preferentially and tightly binds to the outer (extracellular) half of the TMD and consequently opens the KcsA channel. We clearly demonstrate that CL opens the inner gate but has little effect on the inactivation gate (Fig. 4D, E, F). The binding of CL to Arg64 and Arg89, both of which face the inter-subunit space, was unequivocally proven in KcsA mutants (Fig. 5). The MD simulations further suggested the formation of CL mediated hydrogen bond bridges between the two Arg residues (Fig. 6). This binding of CL likely shifts the “open-close” conformational transition of the remote inner gate toward the open state, which could be considered an allosteric modulatory effect (Fig. 7). Hence, we discovered a novel type of activation mechanism for the KcsA channel, which is mediated not via the well-known anionic-lipid sensor M0-helix but by interaction with the inter-subunit positively charged residues at the extracellular surface of the KcsA channel. It should be emphasized that long-range allosteric effects are prevalent in various types of proteins, but have not been found for channel–lipid interactions.

The effect of CL, a major component of the *Streptomyces* membrane, on KcsA was not fully elucidated until this study, even though KcsA is one of the most intensively studied ion channels to date. In this regard, the methodology for analyzing MP–lipid interactions is indispensable for the elucidation of the physiological functions of lipids. The procedure adopted in this study, namely identification of MP-bound lipids by the SAMPLIA method and subsequent functional analysis of the lipids, paves the way for a comprehensive investigation of the biological activity of diverse membrane lipids, thereby accelerating the chemico-biological and physiological studies of lipids.

## Materials and Methods

### Reagents

All lipids used in this study were purchased from Avanti Polar Lipids (Alabaster, AL, USA). The sensor chips for the SPR measurements were purchased from GE Healthcare (Little Chalfont, Buckinghamshire, UK). Any other reagents and materials were all purchased from FUJIFILM Wako Pure Chemical Corp. (Osaka, Japan), Tokyo Chemical Inc. (Tokyo, Japan), or Sigma-Aldrich (St. Louis, MO, USA). Expression and purification of KcsA channels were carried out as previous reported (36). M0 mimicking peptide was purchased from GenScript (Piscataway, NJ, USA).

### Preparation of SPR sensor chip and Immobilization of KcsA

The preparation of the sensor chip was made as previously reported (32). wt-KcsA or ΔM0-KcsA solubilized with 0.06% n-dodecyl-β-D-maltoside (DDM) was immobilized onto SAM-modified sensor chip, according to a standard amino coupling method. HBS-EP buffer (10 mM HEPES [pH 7.4], 150 mM NaCl, 3 mM EDTA, 0.05 % (v/v) Tween 20) was used as the running buffer. Briefly, COOH groups on the surface were activated by injecting a mixture of 390 mM 1-(3-dimethylaminopropyl)-3-ethylcarbodiimide (EDC) and 100 mM N-hydrosuccinimide (NHS) for 7 min, followed by the immobilization of KcsA by injecting the solubilized KcsA solution (100 μg mL^−1^) for 20 min. Unreacted carboxyl groups were deactivated by injecting the blocking solution (1 M ethanolamine-HCl [pH 8.5], 0.05 % (v/v) Tween 20 for 7 min, at a flow rate of 5 μL min^−1^.

### KcsA-lipid interaction analysis

SPR analysis were carried out at 25.0 °C using Biacore T100 system (GE Healthcare, Chicago, IL, USA). In all protocols, acidic buffer (10 mM succinic acid [pH 4.0], 200 mM KCl, 3 mM EDTA, 0.05% (v/v) Tween 20) and neutral buffer (10 mM HEPES [pH 7.5], 200 mM KCl, 3 mM EDTA, 0.05% (v/v) Tween 20) were used as running buffers. Evaluation of KcsA-lipid interactions was made within 2 days after KcsA immobilization. Analyte samples were prepared by dissolving dried lipids into the buffer at the concentrations ranging from 10 to 50 μM, and the samples were sonicated prior to use. All procedures were automated, using repetitive cycles of sample injection and regeneration. The binding assays were performed after three cycles of start-up injections to normalize the two flow cells, and just before the injection of lipid solutions, the buffer alone was injected in the first two cycles in order to obtain baseline value. Afterward, five solutions with increasing lipid concentrations were injected for 180 s at a flow rate of 20 μL min^−1^, followed by 180 s of dissociation at the same flow rate. The surface of the chip was regenerated by one sequential injection of regeneration solution (10 mM Gly-HCl [pH 2.5], 0.05% (v/v) Tween 20) for 30 s at a flow rate of 20 μL min^−1^. Association and dissociation of lipid molecules were expressed as sensorgrams, representing the time-dependent changes. To remove the contribution from non-specific binding between SAM and lipids, a blank channel without KcsA immobilization was used as a reference, and the response of the reference channel (SAM-lipid interactions) was subtracted from that of the sample channel. The evaluation of interactions was performed using BiaEvaluation software (GE Healthcare), and kinetic parameters were extracted by a local fit of the corrected sensorgrams using a 1:1 interaction (Langmuir interaction). The correlations of the fitting were evaluated using χ^2^ analyses and residual plots.

### Single-channel recordings

The contact bubble bilayer (CBB) method was applied for the single-channel recordings, as described in the previous report (28). Prior to the experiment, purified KcsA channel was reconstituted into liposomes by dilution as follows. First, pre-prepared POPC or POPC/anionic lipid (3/1, w/w) liposomes were suspended in 200 mM KCl solution, yielding a 2 mg/ml liposome solution. Then, an aliquot of purified KcsA solution containing 0.06% DDM was diluted 50 times with the liposome solution, obtaining the KcsA-reconstituted proteoliposomes with a lipid/protein ratio of 2000 (w/w). Just before the single-channel recording, a small amount of concentrated buffer (succinic acid for pH4 or HEPES for pH7.5) was added to the proteoliposome solution to adjust the pH (final buffer concentration was10 mM). The proteoliposome solutions of different pHs were filled into separate bubble-forming pipettes (tip diameter, approximately 50 μm). The tip of the pipette was immersed in hexadecane in the chamber while observing with an inverted microscope (IX73; Olympus). A small water bubble (diameter, approximately 100 μm) was inflated at the tip of the pipette by pushing the proteoliposome solution out of the pipette into the hexadecane. A lipid monolayer forms spontaneously at the water-hexadecane interface of the bubble. Finally, a lipid bilayer (i.e., CBB) was prepared by contacting two water bubbles lined with a lipid monolayer (Fig. 4a). The KcsA channel spontaneously transfers from the liposomal membrane to the CBB. The channel insertion into the CBB was detected as the appearance of the single-channel current with the membrane potential applied. Since the KcsA channel is activated when the pH of the intracellular side becomes acidic, only the channel molecule whose cytoplasmic domain faces the acidic bubble contributes to the single-channel under asymmetric pH conditions (pH 4/pH 7.5). Thus, the acidic and neutral bubbles can be defined as the intracellular (in) and extracellular (out) sides, respectively, in the CBB experiments. Asymmetric membranes with different lipid compositions in the inner and outer leaflets were prepared by using different lipid compositions in both bubbles (Fig. 4b). The single-channel current was recorded using a patch-clamp amplifier (Axopatch 200B, Molecular Devices, San Jose, CA). The data was passed through a low-pass filter (1 kHz cutoff frequency) and stored in PC using an analog-to-digital converter (5 kHz sampling; Digidata 1550A; Molecular Devices) and pCLAMP software (Molecular Devices).

### Molecular dynamics simulations

For the structure sampling of open state KcsA, we performed MD simulations under the isothermal-isobaric (NPT) ensemble. The Charmm-GUI web-service were used for construct the initial structure of simulated system including KcsA, lipid bilayer and 0.2 M KCl aqueous solution (47). The X-ray crystal structure of the KcsA (PDB ID: 6by3) was employed for the initial geometry of protein. Glu71s were assumed to be protonated. The KcsA and lipid bilayer were embedded in a rectangular water box of about 145×145×95 Å^3^ with a periodic boundary condition. The extracellular side membrane is constructed by PC and anionic lipids whereas the intracellular side membrane consisted of PC only. We performed three individual simulation with different anionic lipids, namely, PA, PG and CL. PC and PA or PG are randomly placed in a 3:1 molar ratio, PC and CL are in a 6:1 molar ratio, but one anionic lipid was placed near each of intermonomer groove. The CHARMM36m parameters were used for protein and lipids, TIP3P was for water (48, 49). The parameters for ions were taken from the previous report (50).

We conducted equilibrium MD simulations for 10 ns with the constrain of distance between amino hydrogen of Arg64 and phosphate oxygen of anionic lipids placed at intermonomer groove, which were followed by 50 ns equilibrium MD simulations without constrain. After the equilibrium MD simulation, 600 ns MD simulations were conducted for the data production. Long-range electrostatic interactions were treated with the particle mesh Ewald method. The SHAKE method was employed for bond constrain for hydrogen atoms. The leapfrog algorithm was used for time integration with an increment of 2 fs at 298.15 K and 1 bar. All the MD simulations were conducted using the AMBER 20 and AmberTools21 program packages (51, 52).

## Acknowledgments

We acknowledge Dr. Misuzu Ueki for preparation of channel proteins, Ms. Masako Takashima and Ms. Masami Miyagoshi for technical assistance in single-channel recordings.

## Author contributions

In.M. carried out SPR analysis. Iw.M. and O.S. prepared KcsA and its mutants, and performed single-channel recordings. Y.N. performed MD simulations. M.N. and Iw.M. supervised the whole study. All authors wrote the manuscript.

## Data and materials availability

All data are available in the main text or the supplementary materials.

