## Supplementary for "Binding of cardiolipin to the KcsA channel at the membrane outer leaflet allosterically opens the inner gate"

**This PDF file includes:**

Supplementary Method
Figures S1 to S4
Tables S1 to S8
References (1 to 2)

**Supplementary Method**

**Preparation of dodecylamine-modified sensor chip.** Modification of chip surface was performed at 25.0 °C using Biacore T100 system, using preparative CM3 sensor chips, purchased from GE Healthcare. In the protocol of this modification, we used HBS-N (10 mM HEPES [pH 7.4], 150 mM NaCl) as a running buffer. Before modification the chip surface was washed with 50 mM NaOH/2-propanol [3/2 (v/v)] for 3 times at a flow rate of 20 µL $min^{-1}$. In the modification of the chip surface, carboxyl groups on the surface were activated by injecting a mixture of 390 mM EDC and 100 mM NHS for 7 min, followed by the immobilization of dodecylamine by injecting the dodecylamine (1.0 mg $mL^{-1}$, 5.4 mM) in 10 mM NaAc solution (pH 5.0, 10 % (v/v) DMSO) for 30 min, and the deactivation of unreacted carboxyl group by injecting 1 M ethanolamine-HCl (pH 8.5) for 7 min, at a flow rate of 5 µL $min^{-1}$.

**Lipid membranes – peptides interaction analysis.** SPR measurement was performed at 25.0 °C using Biacore T100 system. DOPC (5.0 mg, 6.4 umol) and 0.71 umol of DOPG, DOPS, DOPA, DOPE and tetraC18:1 CL were mixed in MeOH/$CHCl_3$, dried under $N_2$ gas flow, and suspended in 1mL of acidic buffer (10 mM succinic acid [pH 4.0], 200 mM KCl, 3 mM EDTA), to prepare multilamellar vesicles (MLVs). The MLV suspension was extruded through 200 nm, and then 100 nm polycarbonate membranes (LiposoFast Liposome Factory, Sigma-Aldrich, St. Louis, MO, USA) (21 passes for each type of membrane), to prepare 100 nm large unilamellar vesicles (LUVs). The LUV solution was then diluted with the buffer used, to prepare 1 mM LUV solution. LUV solution was injected for 40 min over flowcell on the dodecylamine-modified sensor chip at 2 µL $min^{-1}$ to immobilize LUV onto the chip surface,[1] and washed with 50 mM NaOH solution for 3 times at a flow rate of 20 µL $min^{-1}$ to remove immobilized LUV. The amount of immobilized LUV was recorded after 10 min of signal stabilization. Afterward, each peptide solution (1 µg $mL^{-1}$) was injected for 300 s at a flow rate of 10 µL $min^{-1}$, followed by 300 s of dissociation at the same flow rate. The surface of the chip was regenerated by two sequential injections of 0.5 % (w/v) SDS for 120 s and 50 mM NaOH/2-propanol [3/2 (v/v)] at a flow rate of 20 µL $min^{-1}$. To remove the contribution from non-specific binding of peptide to the chip surface, a blank channel without liposome was used as a reference, and the response of the reference channel was subtracted from that of the sample channel. The procedure to calculate the affinity of peptide to lipid membrane was the same as that used for the KcsA-lipid molecules measurements (1).

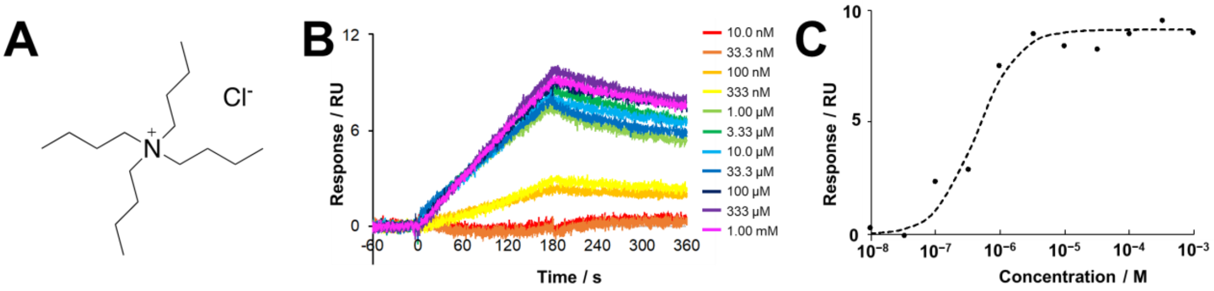


**Figure S1. SPR analysis of the interactions between injected tetrabutylammonium chloride (TBACl) and wt-KcsA immobilized on the $C_6$-SAM modified sensor chip**. **(A)** Structure of TBACl. **(B)** 10 concentrations of TBA were injected in the range of 10 nM to 33 mM dissolved in acidic buffer [10 mM succinic acid, pH 4.0, 200 mM KCl, 3 mM EDTA, 0.05 %(v/v) Tween 20] at a flow rate of 20 μL min$^{-1}$, to obtain a set of sensorgrams. **(C)** The affinity of TBA towards wt-KcsA was assessed from the concentration dependence of the response at 180 s by a fit of Langmuir isotherm binding model. The affinity was calculated as 488 ± 60 nM, which is coincident with the previous channel recording (2).

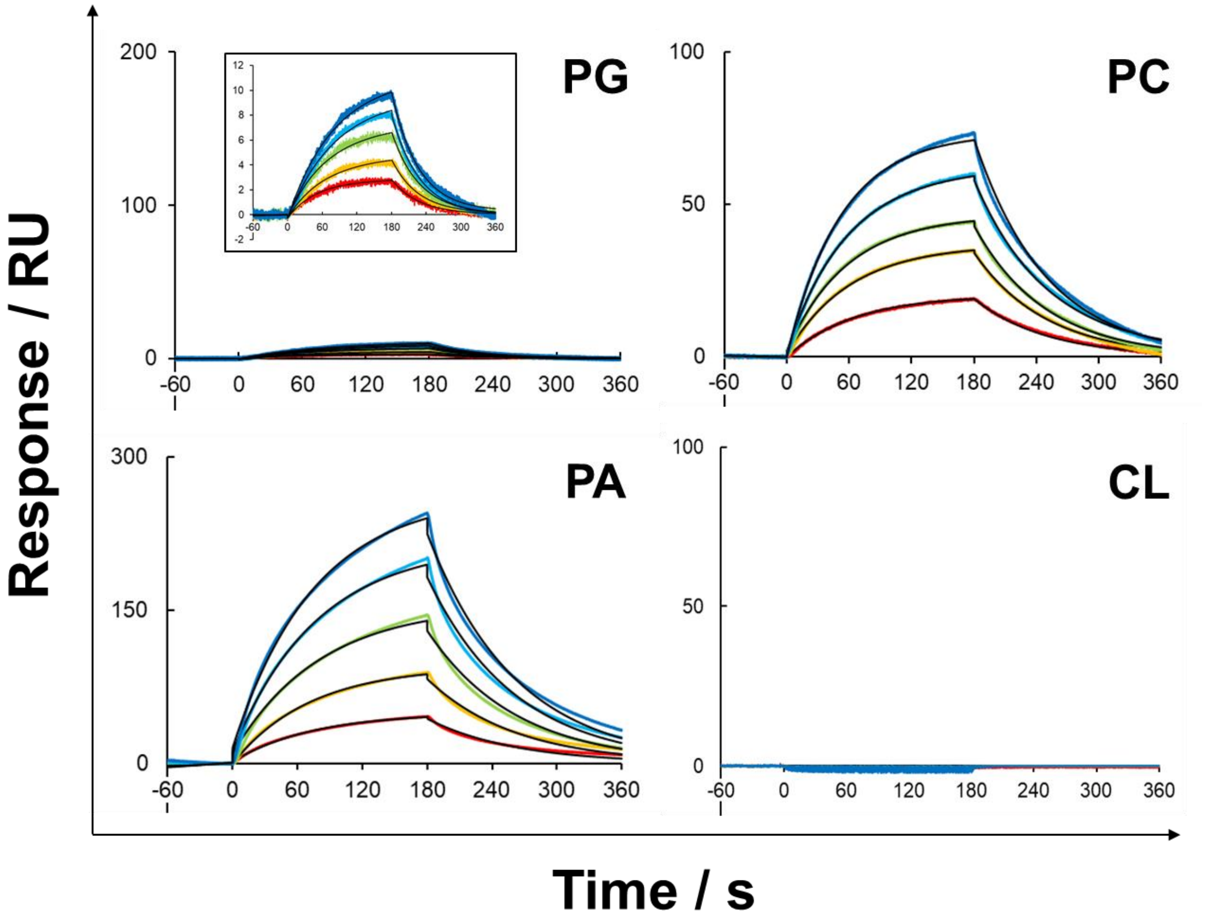


**Figure S2. SPR sensorgrams showing the interactions of PG, PC, PA and CL with wt-KcsA immobilized on the $C_6$-SAM modified sensor chip, at pH 7.5.** 10 (bottom), 20, 30, 40, and 50 (top) μM of the lipids dissolved in neutral buffer [10 mM HEPES, pH 7.5, 200 mM KCl, 3 mM EDTA] were injected at a flow rate of 20 μL min$^{-1}$. Black lines indicate experimentally obtained sensorgrams, while red lines indicate theoretical curves (1:2 heterogeneous ligand binding model).

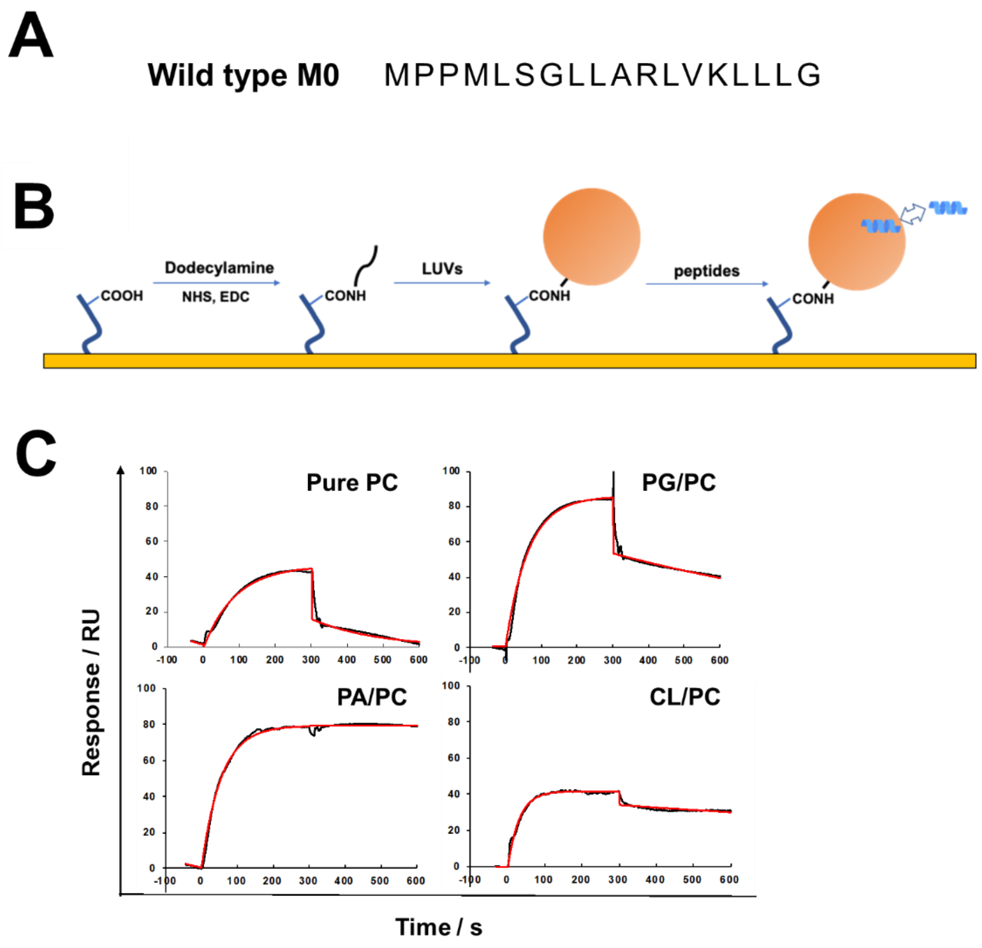


**Figure S3**. **Analysis of the interactions of M0 peptide with liposomes immobilized on the dodecylamine-modified CM3 sensor chip**. **(A)** Amino acid sequence of wild type M0 peptide. **(B)** The procedure of analyzing the interaction. Dodecylamine was coupled with COOH group on the CM3 sensor chip via a conventional amine coupling method. Next, 100 nm LUV solution was injected to be immobilized onto the chip surface. Then the peptide aqueous solution was added to the immobilized liposomes and their interactions were analyzed. **(C)** The sensorgrams showing the interaction of M0 peptide to pure PC, PG/PC, PA/PC and CL/PC membrane in acidic buffer (10 mM succinic acid [pH 4.0], 200 mM KCl, 3 mM EDTA). Black lines indicate experimentally obtained sensorgrams, while red lines indicate theoretical curves (1:1 Langmuir model). The affinity of M0 peptide towards each lipid membrane is listed in Table S3.

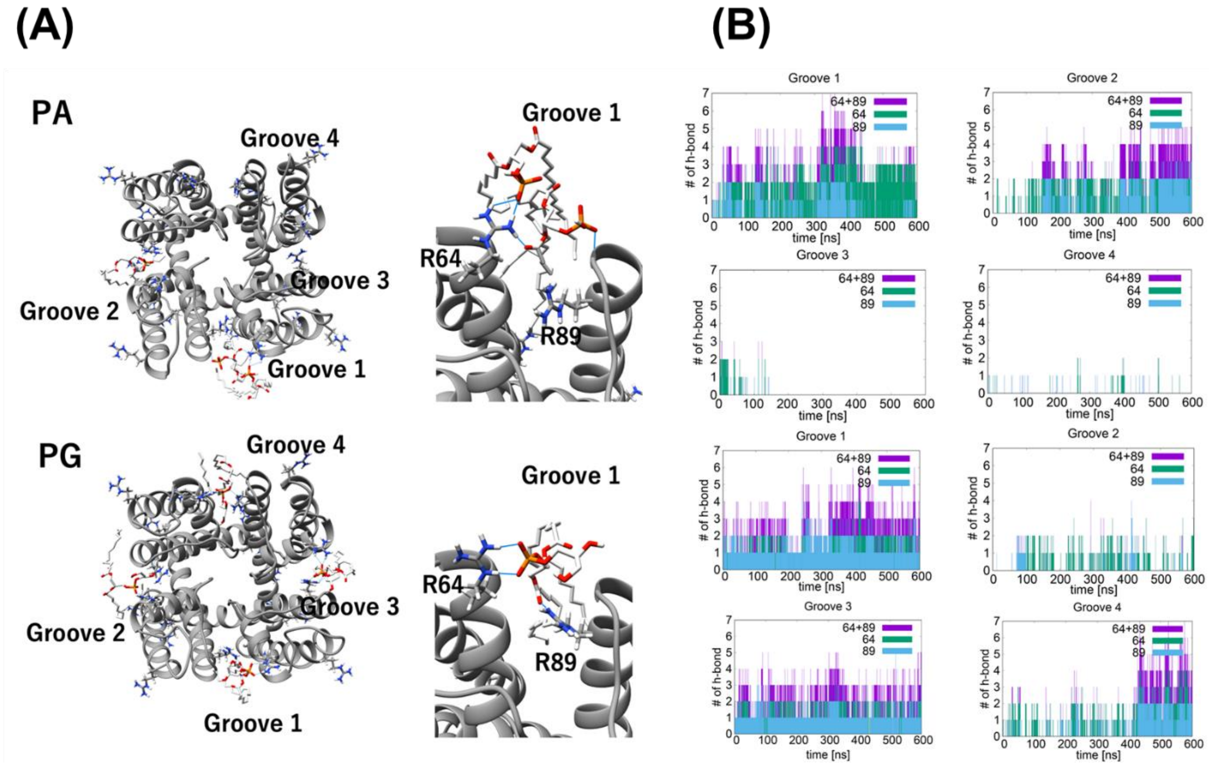


**Figure S4. MD simulations for the binding of PA and PG to KcsA. (A)** Final snapshot of KcsA with 1st neighbor to Arg64 and 89, and their close-up view of the groove 1. Blue lines indicate the possible hydrogen-bond between PA/PG and Arg64 or 89. **(B)** The number of hydrogen bond between PA/PG and Arg64 or 89.

**Table S1**. Particle sizes of lipids in Tween 20-containing buffer, as detected by dynamic light scattering measurement.

| Lipids | Acidic buffer [a] / nm | Neutral buffer [b] / nm |
|---|---|---|
| PC | 10.1 ± 1.8 | 10.0 ± 2.5 |
| PG | 6.2 ± 2.1 | 10.3 ± 2.2 |
| CL | 7.9 ± 1.6 | 11.2 ± 5.1 |
| PA | 8.4 ± 2.8 | 11.0 ± 3.1 |

[a] 10 mM succinic acid [pH 4.0], 200 mM KCl, 3 mM EDTA, 0.05% (v/v) Tween 20

[b] 10 mM HEPES [pH 7.5], 200 mM KCl, 3 mM EDTA, 0.05% (v/v) Tween 20

**Table S2.** Affinities $K_A$ and theoretical maximum response $R_{max}$ of lipids toward wt-KcsA, as determined using SPR method [a]

| Lipid | pH 4.0 (active state) | | pH 7.5 (resting state) | |
|---|---|---|---|---|
| | $K_A$ | $R_{max}$ | $K_A$ | $R_{max}$ |
| PG | $(1.43 \pm 0.13) \times 10^5$ | 249.8 ± 33.7 | $(8.31 \pm 0.56) \times 10^3$ | 383.2 ± 28.1 |
| | $(1.41 \pm 0.19) \times 10^5$ | 17.9 ± 11.5 | $(3.11 \pm 0.73) \times 10^3$ | 1.2 ± 0.6 |
| PC | $(9.91 \pm 0.73) \times 10^3$ | 162.4 ± 7.3 | $(6.55 \pm 0.61) \times 10^3$ | 290.3 ± 15.9 |
| | $(1.13 \pm 0.11) \times 10^3$ | 152.5 ± 21.7 | $(1.82 \pm 0.34) \times 10^3$ | 1.0 ± 0.2 |
| PA | $(8.24 \pm 0.51) \times 10^4$ | 489.8 ± 20.8 | $(5.96 \pm 0.78) \times 10^3$ | 4217.2 ± 560.8 |
| | $(3.16 \pm 0.32) \times 10^3$ | 25.4 ± 5.6 | $(2.57 \pm 1.37) \times 10^3$ | 54.2 ± 22.5 |
| CL | $(6.87 \pm 0.56) \times 10^6$ | 74.1 ± 15.3 | Not detected | |
| | $(3.05 \pm 0.23) \times 10^5$ | 5.5 ± 4.1 | | |

[a] Data are presented as mean ± SD (***N*** = 3). The affinity is shown as the binding constants $K_A$ ($M^{-1}$). Each value was calculated from the sensorgram by fitting to a 1:2 heterogeneous ligand model.

**Table S3.** Affinities $K_A$ and theoretical maximum response $R_{max}$ of lipids toward ΔM0-KcsA at pH 4.0, as determined using SPR method [a]

| Lipid | $K_A$ | $R_{max}$ |
|---|---|---|
| PG | $(4.61 \pm 0.22) \times 10^3$ | $667.4 \pm 25.3$ |
| | $(1.23 \pm 0.61) \times 10^3$ | $2.5 \pm 1.5$ |
| PA | $(3.83 \pm 0.30) \times 10^3$ | $288.7 \pm 29.8$ |
| | $(1.13 \pm 0.23) \times 10^3$ | $45.3 \pm 4.5$ |
| CL | $(4.39 \pm 0.32) \times 10^6$ | $58.3 \pm 7.2$ |
| | $(4.50 \pm 0.37) \times 10^5$ | $8.5 \pm 4.9$ |

[a] Data are presented as mean ± SD (***N*** = 3). The affinity is shown as the binding constants $K_A$ ($M^{-1}$). Each value was calculated from the sensorgram by fitting to a 1:2 heterogeneous ligand model.

**Table S4**. Affinities of M0 peptide towards liposomes at pH 4.0, as determined using SPR [a]

| Liposomes | Affinity $K_A$ / $M^{-1}$ |
|---|---|
| PC | $(3.70 \pm 0.17) \times 10^5$ |
| PG/PC[b] | $(3.60 \pm 0.22) \times 10^7$ |
| PA/PC[b] | $(1.52 \pm 0.13) \times 10^{10}$ |
| CL/PC[b] | $(8.15 \pm 0.59) \times 10^8$ |

[a] Data are presented as mean ± SD (***N*** = 3). [b] The molar ratio was 10/90.

**Table S5**. Open propbability ($P_{open}$) of wt-KcsA channels in CBB [a]

| System | Lipid [b] | $P_{open}$ | SE [c] |
|---|---|---|---|
| In/Out | PG | 0.167 | 0.040 |
| | PA | 0.174 | 0.044 |
| | CL | 0.304 | 0.021 |
| | Cont. (PC) | 0.013 | 0.012 |
| In | PG | 0.165 | 0.031 |
| | PA | 0.191 | 0.024 |
| | CL | 0.255 | 0.017 |
| Out | PG | 0.001 | 0.000 |
| | PA | 0.003 | 0.001 |
| | CL | 0.225 | 0.035 |

[a] A symmetric or asymmetric lipid bilayer was formed by the contact of two bubbles (see Figure 4a, b). [b] These anionic lipids are mixed with PC at an anionic lipid/PC weight ratio of 1/3. [c] ***N*** = 3-14.

**Table S6**. Open propbability ($P_{open}$) of E71A mutant in CL-containing CBB [a]

| System [b] | $P_{open}$ | SE [c] |
|---|---|---|
| In/Out | 0.984 | 0.009 |
| In | 0.945 | 0.027 |
| Out | 0.852 | 0.050 |
| Cont.(PC) | 0.067 | 0.027 |

[a] CL was mixed with PC at a CL/PC weight ratio of 1/3. [b]A symmetric or asymmetric lipid bilayer was formed by the contact of two bubbles (see Figure 4a, b). [c] $\boldsymbol{N}$ = 4-8.

**Table S7.** Affinities $K_A$ and theoretical maximum response $R_{max}$ of CL toward Arg mutants of KcsA, as determined using SPR method [a]

| Mutation | Affinity $K_A$ / $M^{-1}$ | Theoretical maximum response $R_{max}$ |
|---|---|---|
| E71A | $(1.61 \pm 0.19) \times 10^6$ | $654.6 \pm 85.8$ |
| | $(7.66 \pm 0.33) \times 10^4$ | $2.8 \pm 1.3$ |
| E71A/R52Q | $(2.57 \pm 0.17) \times 10^6$ | $533.4 \pm 33.9$ |
| | $(3.05 \pm 0.18) \times 10^5$ | $23.8 \pm 10.3$ |
| E71A/R64Q | $(4.56 \pm 0.33) \times 10^4$ | $1147.0 \pm 37.7$ |
| | $(2.27 \pm 0.45) \times 10^4$ | $0.2 \pm 0.1$ |
| E71A/R89Q | $(3.03 \pm 0.27) \times 10^4$ | $1518.4 \pm 75.2$ |
| | $(1.01 \pm 0.10) \times 10^4$ | $0.1 \pm 0.1$ |
| E71A/R64Q/R89Q | $(3.26 \pm 0.36) \times 10^4$ | $1253.4 \pm 57.0$ |
| | $(1.35 \pm 0.19) \times 10^4$ | $0.1 \pm 0.1$ |

[a] Data are presented as mean ± SD ($N = 3$). Each value was calculated from the sensorgram by fitting to a 1:2 heterogeneous ligand model.

**Table S8.** Open probability ($P_{open}$) of Arg mutants of KcsA in CL-containing CBB [a]

| Mutation | System [b] | $P_{open}$ | SE [c] |
|---|---|---|---|
| E71A/R52Q | In/Out | 0.943 | 0.019 |
| | In | 0.976 | 0.009 |
| | Out | 0.931 | 0.021 |
| E71A/R64Q | In/Out | 0.954 | 0.029 |
| | In | 0.886 | 0.080 |
| | Out | 0.061 | 0.025 |
| E71A/R89Q | In/Out | 0.917 | 0.042 |
| | In | 0.993 | 0.002 |
| | Out | 0.045 | 0.014 |
| E71A/R64Q/R89Q | In/Out | 0.974 | 0.007 |
| | In | 0.764 | 0.080 |
| | Out | 0.065 | 0.032 |

[a] CL was mixed with PC at a CL/PC weight ratio of 1/3. [b] A symmetric or asymmetric lipid bilayer was formed by the contact of two bubbles (see Figure 4a, b). [c] ***N*** = 3-19.
